# Brain Representations of Natural Sound Statistics

**DOI:** 10.1101/2025.06.17.660096

**Authors:** Yousef Mohammadi, Alexander J. Billig, Joel I. Berger, Timothy D. Griffiths

**Affiliations:** Department of Imaging Neuroscience, University College London 12 Queen Square, London, WC1N 3AR, United Kingdom; Newcastle University Medical School Framlington Place, Newcastle upon Tyne, NE2 4HH, United Kingdom; UCL Ear Institute, University College London 332 Grays Inn Road, London, WC1X 8EE, United Kingdom; Department of Neurosurgery, The University of Iowa Iowa City, IA, 52242, United States of America

## Abstract

Natural sound textures (e.g., rain, crackling fire) are perceptually defined by time-averaged summary statistics. While previous studies have examined neural responses to natural sounds, little is known regarding the neural processing of the statistics underlying these sounds. To study neuronal correlates of these statistics, we measured brain responses to synthetic sound textures in which statistical structure was systematically varied while preserving the texture category. Using two fMRI experiments (males and females), we examined neural responses along the ascending auditory pathway, within auditory cortex and medial temporal lobe (MTL) regions implicated in pattern analysis. In Experiment 1, we parametrically varied the full set of texture statistics, creating sounds with different levels of naturalness. In Experiment 2, we selectively manipulated high-level statistics (cochlear skewness and kurtosis, cochlear and modulation correlations) while holding low-level statistics (cochlear mean and modulation power) constant. Increasing texture naturalness produced graded increases in BOLD responses across bilateral primary and nonprimary auditory cortex in both experiments, although overall responses were weaker in Experiment 2. This reduction suggests that low-level statistics contribute substantially to response magnitude, even though higher-order statistics are sufficient to elicit graded responses. We also observed modulation in MTL regions, including entorhinal cortex, in Experiment 1. Moreover, functional connectivity between hippocampus and auditory cortex increased for more degraded (less natural) textures, suggesting a modulatory rather than representational role for MTL in texture processing. Together, these findings show that sensitivity to texture statistics is distributed across the auditory cortex and highlight MTL-auditory interactions when texture structure is ambiguous.

**Significant Statement:** Natural sound textures such as rain or crackling fire are perceptually defined by time-averaged summary statistics that support efficient auditory perception, yet how the human brain represents these statistics remains unclear. Using fMRI and synthetic sound textures in which statistical structure was systematically manipulated, we found that both primary and nonprimary auditory cortex were sensitive to texture statistics, exhibiting a graded and distributed representation of these acoustic features. We also observed increased functional connectivity between the hippocampus and auditory regions when texture structure was degraded or the sounds were unnatural. Together, these results indicate that sound texture statistics are encoded across multiple levels of auditory cortex and further suggest a modulatory role for hippocampus under conditions of heightened perceptual uncertainty.

## Introduction

Everyday listening environments are filled with rich acoustic backgrounds, such as rustling leaves, falling rain, or crackling fire. There is growing evidence that the auditory system has evolved to process behaviourally relevant natural sounds (Theunissen and Elie, 2014), and the broader idea that sensory systems adapt to the statistical structure of the natural environment has a long history (Barlow, 1961). Early work on statistical regularities in natural sounds includes the research of Voss and Clarke (1975), who demonstrated that pitch and amplitude fluctuations in extended segments of speech and music exhibit a 1/f distribution, as well as evidence for power-law amplitude distributions (Attias and Schreiner, 1997). Neurons across the auditory pathway preferentially encode such statistics (Garcia-Lazaro et al., 2006, 2011). Singh and Theunissen (2003) further described the low-pass nature of spectral and temporal modulations in natural sounds, corroborating and extending the findings of Voss and Clarke (1975). Despite this history, the literature on the relevance of natural sound statistics to auditory processing and perception remains relatively sparse. In this context, the study by McDermott and Simoncelli (2011) offered a significant advance by showing that imposing a specific set of statistical parameters, extracted from spectral and modulation-based representations, on white noise can produce highly realistic synthetic textures, suggesting that these parameters act as “summary statistics” for describing sound textures.

Different natural sounds have distinct statistical properties (McDermott and Simoncelli, 2011). This is shown in detail by Mishra et al (2021), who examined sound texture summary statistics by decomposing them into a smaller set of components. For instance, marginal moments distinguish “sparse” or “bursty” textures, characterized by intermittent energy bursts, from more stationary textures. Cochlear correlations differentiate “highly correlated” textures (e.g., applause) from “poorly correlated” ones (e.g., water-like textures). Modulation power distinguishes rapidly modulating sounds (e.g., fast wing flaps of a bee) from slowly modulating ones such as ocean waves, whereas modulation correlations differentiate textures exhibiting sudden phase transitions or onset-offset patterns (e.g., firecrackers). Together, these findings suggest that different texture types depend on different subsets of summary statistics, supporting a distributed representation of sound textures (Santoro et al., 2014). Furthermore, these statistics can be viewed as forming a hierarchical processing framework: low-level features like marginal moments can be derived from activity in individual cochlear channels, while high-level features like cochlear correlations require integration across frequency channels along the tonotopic axis. This indicates that higher-level statistics should be represented in higher-order cortical areas that reflect hierarchical organization (McDermott and Simoncelli, 2011; Peng et al., 2024).

Here, we investigated cortical responses to sound textures with varying levels of summary statistics. We conducted two experiments. In Experiment 1, we generated synthesized texture sounds in which all statistics were manipulated - low-level (cochlear mean and power, modulation power) and high-level (cochlear skewness and kurtosis, cochlear and modulation correlations). This manipulation produced four naturalness levels (0.25, 0.4, 0.63, and 1), corresponding to different proportions of texture–noise statistics. In Experiment 2, to isolate the contribution of higher-order statistics, only the high-level statistics were varied across naturalness levels, while all low-level statistics were held constant. Based on hierarchical models of auditory processing, we predicted that primary auditory cortex would show reduced BOLD responses in Experiment 2 relative to Experiment 1, whereas responses in nonprimary auditory regions would be preserved.

Across both experiments, we found that synthesized textures with higher naturalness elicited stronger BOLD responses in primary and nonprimary auditory regions. In Experiment 2, where low-level modulation power was held constant, overall response amplitudes in both regions were weaker than in Experiment 1. Together, these results indicate a distributed cortical representation of both low- and high-level texture statistics. We also observed that functional connectivity between auditory regions and the hippocampus was modulated by texture naturalness, with increased connectivity for unnatural textures. This pattern suggests a potential top-down modulatory role for the hippocampus, while the core representation of texture statistics resides in auditory cortex.

## Methods

In both experiments the same procedure, acquisition, and analysis pipeline were used. The only difference was in how acoustic features of the stimuli were manipulated, as outlined below.

### Participants

For Experiment 1, thirty individuals (aged 18-40 years, median 26 years, 10 males, 20 females) and for Experiment 2, fifteen individuals (aged 19-33 years, median 24 years, 7 males, 8 females) with self-reported normal hearing took part. The sample size for Experiment 2 was determined using a power calculation based on the results of Experiment 1 (see Univariate fMRI Analysis section). Informed consent was obtained from all participants, who were compensated for their time. The study was approved by the University College London Research Ethics Committee.

### Stimuli

Sound texture stimuli were synthesized based on an auditory texture model (McDermott and Simoncelli, 2011). Texture statistics (*ξ_texture_*) were first measured in 7-s original recordings of real-world sound textures (“Birds,” “Rain,” “Fire,” and “Insects”). These sounds were chosen as they have distinct statistical structure. The statistics include marginal moments of each cochlear envelope (the mean, coefficient of variation, skewness and kurtosis), the pairwise correlation coefficient between cochlear envelopes, the power in each modulation band, and pairwise correlations across modulation bands (Figure 1). With the same procedure, statistics (*ξ_noise_*) of 7-s Gaussian noise were also calculated. Texture morphs with different level of texture statistics were generated by synthesizing a texture from a point along a line in the space of statistics between the original texture and noise (*ξ_synthesis_* Δ*ξ_texture_* + (1 - Δ)*ξ_noise_*), using 3-s samples of Gaussian noise as a seed (McWalter and McDermott, 2018). Four logarithmic-scaled levels of Δ were selected (0.25, 0.4, 0.63, 1). In Experiment 1, we matched for cochlear envelope means along Δ levels; in other words, the cochlear envelope means in the noise statistics, *ξ_noise_*, were replaced with those of the texture statistics, *ξ_texture_*, so the *ξ_synthesis_* at each Δ level had similar cochlear means (Supplementary Figure S1). In Experiment 2, in addition to matching cochlear envelope means, we also matched cochlear envelope variance and modulation power across conditions (i.e., the envelope means, variance, and modulation power in the noise statistics, *ξ_noise_*, were replaced with those of the texture statistics, *ξ_texture_*) (Figure 1). To quantitatively illustrate how the statistics vary across Δ levels, we computed, for each texture, the Euclidean distance between the statistics at Δ = 0.4, 0.63, and 1 and those at Δ = 0.25. This procedure was repeated across multiple synthesized exemplars, and the resulting distance measures were averaged. The results, shown in Supplementary Figure S2, confirm that in Experiment 2 the modulation power statistics remain effectively constant across Δ values.

**Figure 1.**
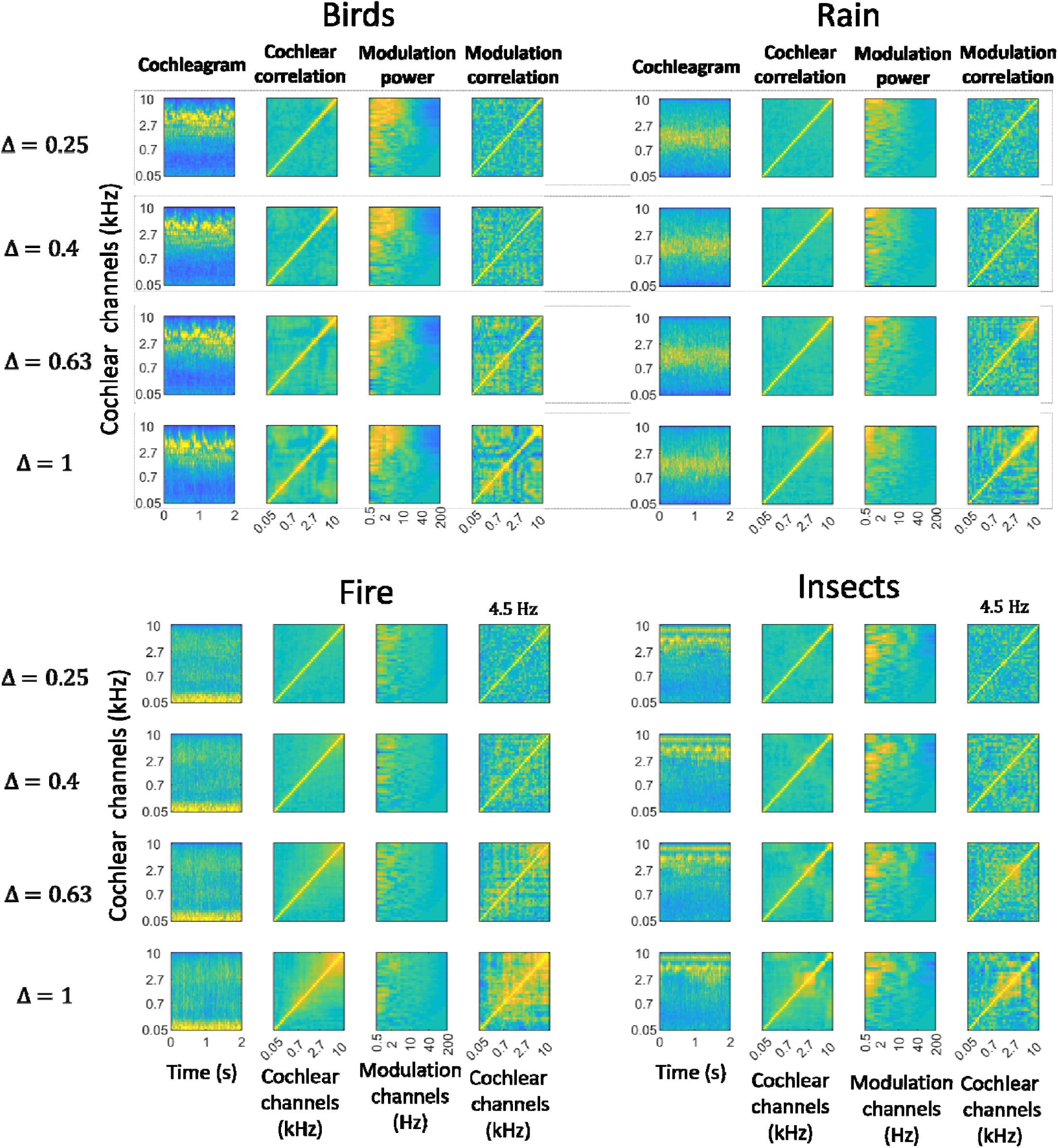
Texture Cochleagram and summary statistics. High-level statistics (cochlear channel kurtosis and skewness, cochlear correlation and modulation correlation) systematically varied with different proportions of texture and noise statistics defined here as = [0.25, 0.4, 0.63, and 1], i.e., higher values denote higher naturalness. for four texture sounds (Birds, Rain, Fire and Insects). Other statistics (cochlear channel mean and variance, and modulation power) were held constant during synthesis.

In both experiments, for each Δ level, several different exemplars were synthesized using different random seeds. All synthesis parameters were set as in (McWalter and McDermott, 2018). The 2-s segments (with random beginning point between 0-1s) were selected from synthesized texture sounds. The root mean square (RMS) of each sound was normalized to a desired level of 0.1, and then the long-term average loudness of each sound was adjusted to 22 sones using the Glasberg and Moore method (Moore et al., 2016). A 10-ms cosine ramp was then applied to the beginning and end of each sound. Sounds were delivered using a bespoke high-fidelity delivery system based on a manifold of parallel piezoelectric transducers. An equalizer filter was applied (prior to RMS normalization) to compensate for sound delivery system attenuation.

### Procedure

The experiment used a sound discrimination task (Figure 2A). Each trial began with a 1-second visual alert of “Sounds to start” displayed on the screen, followed by the presentation of the reference and test sounds with an interstimulus interval of 2-4 s while fixation cross displayed on the screen. After the test sound, a message “Same or different?” appeared on screen. In each trial, reference and test sounds had the same texture type (e.g., birds) and only differed in their level of Δ. Participants were instructed to respond as quickly as possible without making mistakes within a 3-second time window, using one of two available buttons on a keypad. They had to decide whether the reference and test sounds were the same or different. Each trial ended with a resting period of variable length, jittered between 2 and 4 s during which a fixation cross was displayed on the screen.

**Figure 2.**
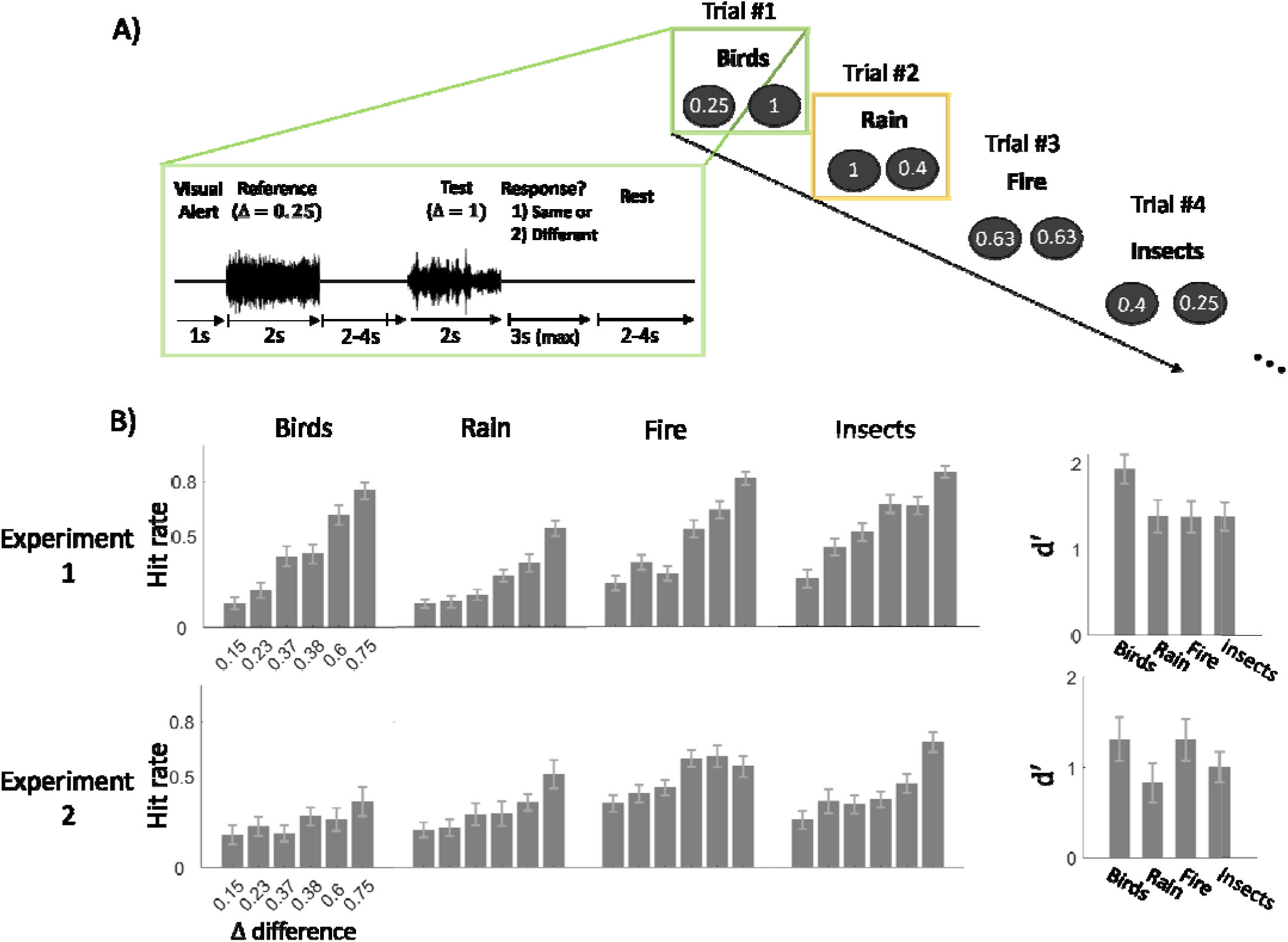
Texture perception paradigm and discrimination performance across participants. A) Each trial consisted of two sounds, a “reference” and a “test” that either shared the same or differed in their level of texture statistics, denoted as Δ: [0.25, 0.4, 0.63, 1] (Δ representing the proportion of texture statistics features over statistics of random noise sound). These sounds were separated by a short interstimulus interval. Four types of natural texture sounds were used: Birds, Rain, Fire, and Insects. B) Discrimination performance (measured as hit rate) is shown for Experiment 1 (top) and Experiment 2 (bottom). Performance is plotted as a function of the absolute difference between the Δ values of the reference and test sounds. Only trials with a nonzero Δ difference (75% of all trials) are included, and the hit rate shows the proportion of trials in which participants successfully detected the difference in Δ levels between the reference and test sounds. The right panel shows the d′ score for each texture sound. In Experiment 1, the cochlear envelope mean was matched for each Δ level and in Experiment 2 cochlear mean, variance, and modulation power were matched. Error bars represent the standard error of the mean.

Trials were presented in 5 runs of about 10 min, each containing 52 trials, totalling 260 trials. Between runs, participants were permitted a short rest but were required to remain still. Four texture sounds (“Birds,” “Rain,” “Fire,” and “Insects”) were used. In each run, trials were divided into 13 groups, each containing all four texture types. Within each group, the order of the textures was randomized.

Different levels Δ were used for the reference and test sounds in 75% of the trials (“different” trials), while identical levels were used in the remaining 25% (“same” trials). The order of different and same trials was randomized. An equal number of each Δ level was utilized in both different and same trials. In different trials, the number of trials with a greater Δ for the reference and test sounds was counterbalanced, and the order of these trials was randomized.

Prior to the main experiment, participants completed a practice session outside the scanner room consisting of 16 trials with different sound exemplars than those used in the main experiment, intended to familiarize them with the task. The experiment was controlled using Psychtoolbox and custom MATLAB code.

### MRI data acquisition

Data were acquired using a 3-Tesla Siemens MAGNETOM Prisma MR scanner (Siemens Healthcare) with a 32-channel receiver head coil at the Department of Imaging Neuroscience (London, UK). Whole-brain T2*-weighted functional images were acquired using 3D echo-planar imaging (EPI) with field of view 192 x 192 x 144 mm, voxel size 3 mm isotropic, 48 partitions, 3D acceleration factor 2, repetition time (TR) 44ms, volume acquisition time 1.056 s, anterior-to-posterior EPI phase encoding, and bandwidth 2298 Hz/Px. Field maps (short echo time (TE) = 10.00 ms, long TE = 12.46 ms) and a whole-brain T1-weighted anatomical image (MPRAGE, field of view 256 x 256 x 176 mm, voxel size 1 mm isotropic, GRAPPA acceleration factor 2, TR = 2530 ms, TE = 3.34 ms) were acquired during the same scanning session.

During scanning sessions peripheral measurements of subject pulse rate and respiratory signals were made together with scanner slice synchronisation pulses using the Spike2 data acquisition system (Cambridge Electronic Design Limited, Cambridge UK). The cardiac pulse signal was measured using an MRI compatible pulse oximeter (Model 8600 F0, Nonin Medical, Inc. Plymouth, MN) attached to the subject’s left index finger. The respiratory signal, thoracic movement, was monitored using a pneumatic belt positioned around the abdomen close to the diaphragm.

A physiological noise model was constructed to account for artifacts related to cardiac and respiratory phase and changes in respiratory volume using an in-house developed Matlab toolbox (Hutton et al., 2011). This resulted in a total of 14 regressors which were sampled at a reference slice in each image volume to give a set of values for each time point. The resulting regressors were included as confounds in the first level analysis for each subject.

To ensure that sounds were presented at a comfortable listening level, we conducted a calibration test inside the MRI scanner with an EPI sequence running, prior to the main experiment. In Experiment 1, participants listened to a continuous sound composed of a mixture of synthesised textures, presented initially at a baseline sound pressure level (SPL) of 68 dB. They were instructed to press a button to gradually increase the volume until they reached a level that they found comfortable, while ensuring that all texture sounds remained clearly audible. The average SPL across participants was 82 dB. In Experiment 2, sounds were initially presented at an SPL level of 80 dB and the final average level across participants was 84 dB.

### MRI preprocessing

MRI data were processed in SPM12 (Department of Imaging Neuroscience, UCL). Each participant’s functional images were unwarped using their field map and realigned to the first volume. Functional and anatomical images were coregistered to the mean functional image. For the univariate analyses, the coregistered images were normalised to the SPM12 template (avg305T1) in MNI space and the functional images were spatially smoothed using with a Gaussian kernel with 8 mm full-width at half-maximum (FWHM).

After the data were preprocessed using SPM, the artifact detection (ART) toolbox, implemented in CONN toolbox (Whitfield-Gabrieli and Nieto-Castanon, 2012), was used to detect outliers from smoothed images. Time points in the series were marked as outliers if the global signal exceeded 3 standard deviations and the movement exceeded 1 mm. Additionally, five covariates generated using the aCmpCorr method (Behzadi et al., 2007), which uses principal component analysis on the measurements made in the white matter (WM) and cerebrospinal fluid (CSF) of each individual subject’s segmented WM and CSF masks, were included as confounds in the first level analysis for each subject.

### Univariate fMRI Analysis

To examine how BOLD activity varied with the Δ (i.e., the proportion of the texture statistics: 0.25, 0.4, 0.63, and 1.0) we conducted a whole brain voxel-wise univariate analysis using a general linear model in SPM12. At the first level, each participant’s time series was modelled with separate regressors for each texture category (Birds, Rain, Fire, and Insects). This allowed us to control for potential category-specific differences while examining Δ effects across all textures. For each regressor, we also included a parametric modulator corresponding to the Δ. All regressors were convolved with the canonical hemodynamic response function. Data were high-pass filtered (cut-off: 1/128 Hz) to remove low-frequency drifts. First-level contrasts of the main effect of the parametric modulation of Δ were estimated for each participant. These contrast images were then entered into second-level group analyses using a random-effects model. To identify regions showing a positive linear relationship with Δ, we conducted a one-sample t-test on the parametric modulation contrast across participants. Small-volume correction was applied within the auditory cortex and the medial temporal lobe (MTL), and because these corrections were performed in two independent regions, the region-wise significance threshold was Bonferroni-adjusted. Anatomical labeling of auditory subregions was based on the Julich-Brain Atlas (Amunts et al., 2020) (Figure 3B).

**Figure 3.**
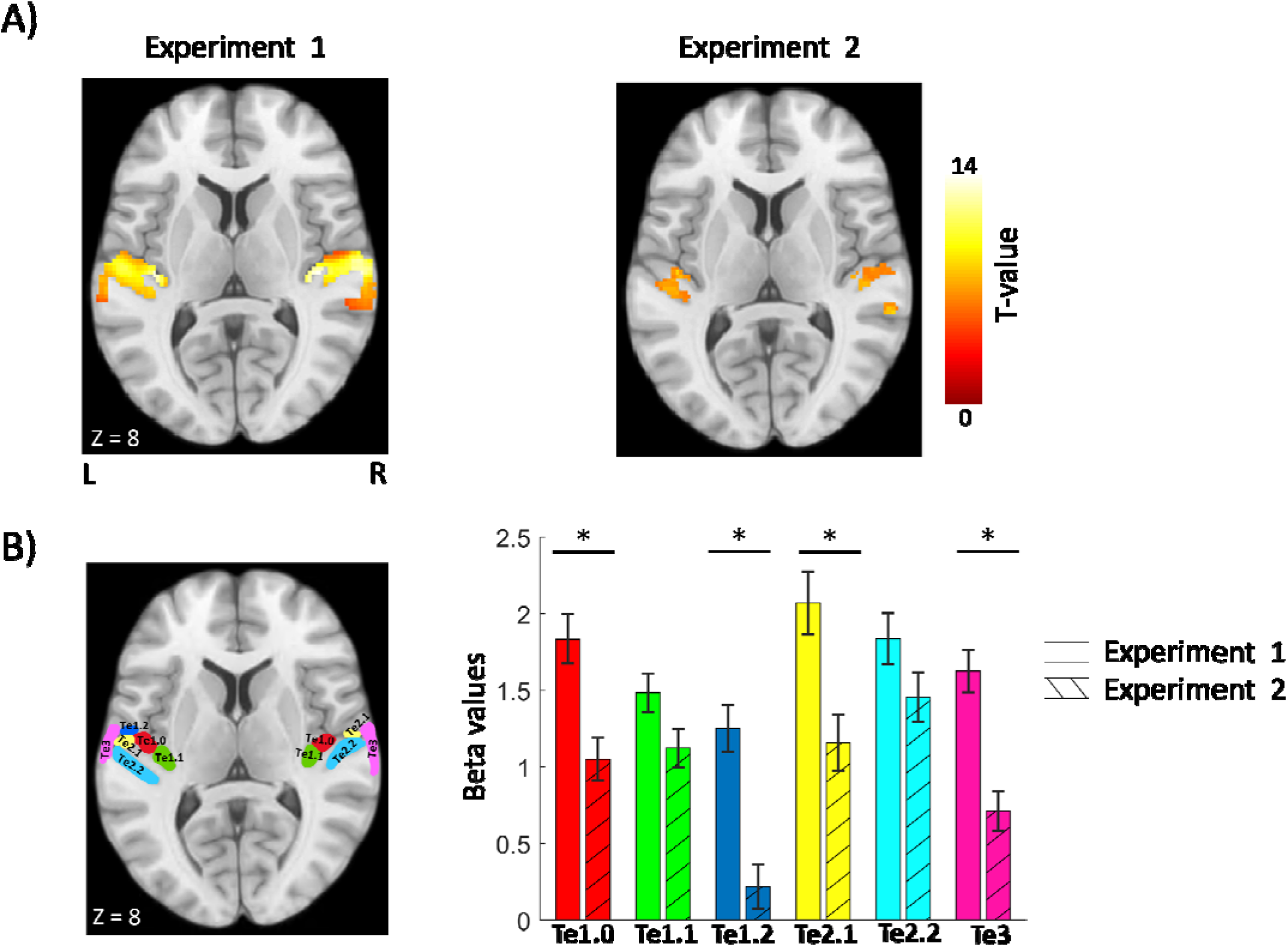
A) Auditory cortical responses to proportional increase in texture statistics during texture perception (at p<0.05, FWE-corrected for auditory cortex). BOLD activity was measured while participants listened to pairs of synthetic textures varying in texture statistics, denoted as Δ: [0.25, 0.4, 0.63, 1] (Δ representing the proportion of texture statistics features over statistics of random noise sound). Bilateral auditory cortex showed increased activation with higher Δ. In Experiment 1, only cochlear envelope mean was matched across Δ level and in Experiment 2, in addition to cochlear mean, cochlear variance and modulation power were also matched. B) Beta values associated with the effect of Δ were extracted for each auditory subregion. Error bars represent the standard error of the mean.

To compare BOLD responses between the two experiments, we conducted a region-of-interest (ROI) analysis in which beta values for each ROI, defined using the Julich-Brain Atlas, were extracted and entered into a mixed-model ANOVA, with experiment as a between-subjects factor and ROI as a within-subjects factor. Post-hoc t-tests were then performed to assess differences between experiments within each ROI. Assumptions of normality and homogeneity of variance, assessed using the Shapiro–Wilk and Bartlett tests, respectively, were not violated; therefore, standard two-sample t-tests were applied. Bonferroni-corrected p-values are reported.

In Experiment 2, we included 15 participants. Because high-level acoustic statistics were not manipulated in this experiment, we expected effects in higher-level auditory regions to resemble those observed in Experiment 1. To estimate the required sample size, we extracted beta values from area Te3 in Experiment 1, which showed a large effect (Cohen’s d = 2.15). An effect size of this magnitude corresponds to an estimated required sample size of n = 7 (α = 0.05, power = 0.95). However, anticipating that the effect size might be reduced in Experiment 2 due to the control of modulation power, we recruited 15 participants to ensure that any observed differences would not be attributable to insufficient statistical power.

Effect sizes in the medial temporal lobe (MTL) were more moderate. For example, in the entorhinal cortex, where we observed a significant effect in Experiment 1, the effect size was Cohen’s d = 0.72, corresponding to an estimated required sample size of n = 28 for comparable power. However, because the primary aim of Experiment 2 was to characterize responses in auditory cortex, we focused on ensuring adequate power for effects in this region and therefore included 15 participants.

### Psychophysiological Interaction (PPI) Analysis

We used the psychophysiological interaction (PPI) method, as implemented in the CONN toolbox, to test whether connectivity between auditory subregions and medial temporal lobe (MTL) areas was modulated by the level of texture naturalness (Δ). We defined seed regions in auditory regions of interest (ROIs; Te1.0, Te1.1, Te1.2, Te2.1, Te2.2, Te3, STS1, STS2) according to the Julich-Brain Atlas (Amunts et al., 2020), and for the MTL, we included the entorhinal cortex, hippocampus, parahippocampal gyrus, and amygdala for each hemisphere. PPI analysis was performed for each ROI pair (ROI-to-ROI analysis) for each subject. The effect of movement and physiological parameters on the BOLD signal was reduced by regressing out motion and physiological artifacts, along with their first-order temporal derivative, and covariates generated using the aCmpCorr method from WM and CSF by running whole-brain voxel wise regression.

Following CONN’s standard procedure for gPPI ROI-to-ROI analysis (bivariate regression), we first applied an ROI-level MVPA omnibus test to all ROIs from both hemispheres (24 ROIs total), restricting the analysis to within-hemisphere connections (e.g., connections between right Te3 and the other 11 right-hemisphere ROIs). This multivariate test evaluates whether each ROI exhibits a significant overall pattern of connectivity differences. The test produced 24 p-values, one per ROI, which were then corrected for multiple comparisons at the ROI level using FDR (p < 0.05). ROIs that survived correction were subsequently examined with post-hoc univariate analyses, in which each individual connection associated with that ROI was tested using a two-tailed one-sample t-test. These post-hoc p-values were further FDR-corrected within each ROI (p < 0.05) to identify the specific connections driving the significant omnibus effect. This procedure allowed us to characterize connectivity patterns between auditory regions and medial temporal lobe structures as a function of increasing or decreasing texture naturalness.

### Code and data availability

All fMRI and behavioural data are available <u>*here*</u>.

## Results

### Behavioural performance

Participants were presented with two sounds separated by a short interval. Each pair consisted of sounds drawn from the same texture but containing either identical or different levels of texture statistics (Figure 2A). Participants were asked to judge whether the two sounds were the same or different. The primary goal of having an active discrimination task inside the scanner was to keep participants engaged in listening and to make sure that differences in texture statistics were detectable inside the MRI scanner. Nevertheless, we compared behavioural performance between experiments by calculating the mean hit rate across texture sounds at each level of Δ difference (i.e., the difference between the reference and test naturalness levels; Figure 2B). A mixed-model ANOVA was conducted with Experiment as the between-subjects factor and Δ-difference (0.15, 0.23, 0.37, 0.38, 0.6, and 0.75) as the within-subjects factor. The analysis revealed a significant Experiment × Δ-difference interaction, F(1, 43) = 7.1, p < 0.001, as well as a main effect of Δ-difference, F(5, 215) = 41, p < 0.001, but no main effect of Experiment, F(1, 12) = 2.5, p = 0.10.

Follow-up comparisons revealed that the two experiments differed only at the largest Δ-difference (0.75): participants in Experiment 1 performed significantly better than those in Experiment 2 (p = 0.001). This result can be explained by the structure of the stimuli at the Δ-difference of 0.75, which corresponds to the contrast between sounds with Δ = 1 and Δ = 0.25. Although the Δ = 1 stimuli were similar across experiments, the Δ = 0.25 stimuli diverged: in Experiment 2, cochlear variance and modulation power were matched to the fully natural texture, whereas in Experiment 1 they were reduced to 25% of the natural value. The performance difference at this level therefore suggests that modulation power, as a low-level statistic, serves as a behaviourally salient cue for judging texture naturalness.

### Neural responses to texture statistics

In Experiment 1, univariate analysis showed significant positive parametric modulation of texture naturalness (i.e., stronger activity for more natural textures) in bilateral primary and nonprimary auditory cortex (p < 0.05, FWE corrected for the volume of auditory cortex), including subregions Te1.0, Te1.1, Te1.2, Te2.1, Te2.2, and Te3 (Figure 3A). No significant negative parametric effects of naturalness (i.e., stronger activity for less natural textures) were observed in auditory cortex. In Experiment 2, a similar pattern of activity in primary and non-primary regions was observed (p < 0.05, FWE corrected for the volume of auditory cortex). The ROI analysis revealed a significant interaction between Experiment and auditory subregion, F(5, 215) = 3.48, p = 0.004, as well as main effects of Experiment, F(1, 43) = 12.6, p = 0.009, and ROI, F(5, 12) = 18.0, p < 0.0001. We then compared responses between the two experiments within each ROI. Greater responses in Experiment 1 were observed in Te1.0 (p = 0.017), Te1.2 (p = 0.0004), Te2.1 (p = 0.037), and Te3 (p = 0.0007). No significant differences emerged in Te1.1 (p = 0.48) or Te2.2 (p = 0.90).

In addition to auditory cortex, in Experiment 1, we observed activity in medial temporal lobe regions, covering entorhinal cortex, at an uncorrected threshold (p < 0.001) (not shown here). We then performed modelling of BOLD responses for reference and test sounds separately, to see whether this activity was present for both sounds. We observed an effect of in entorhinal cortex (EC) (p = 0.01, FWE-corrected for MTL volume) (Figure 4) for the test sound presentation. No significant effects of were observed in MTL in Experiment 2.

**Figure 4.**
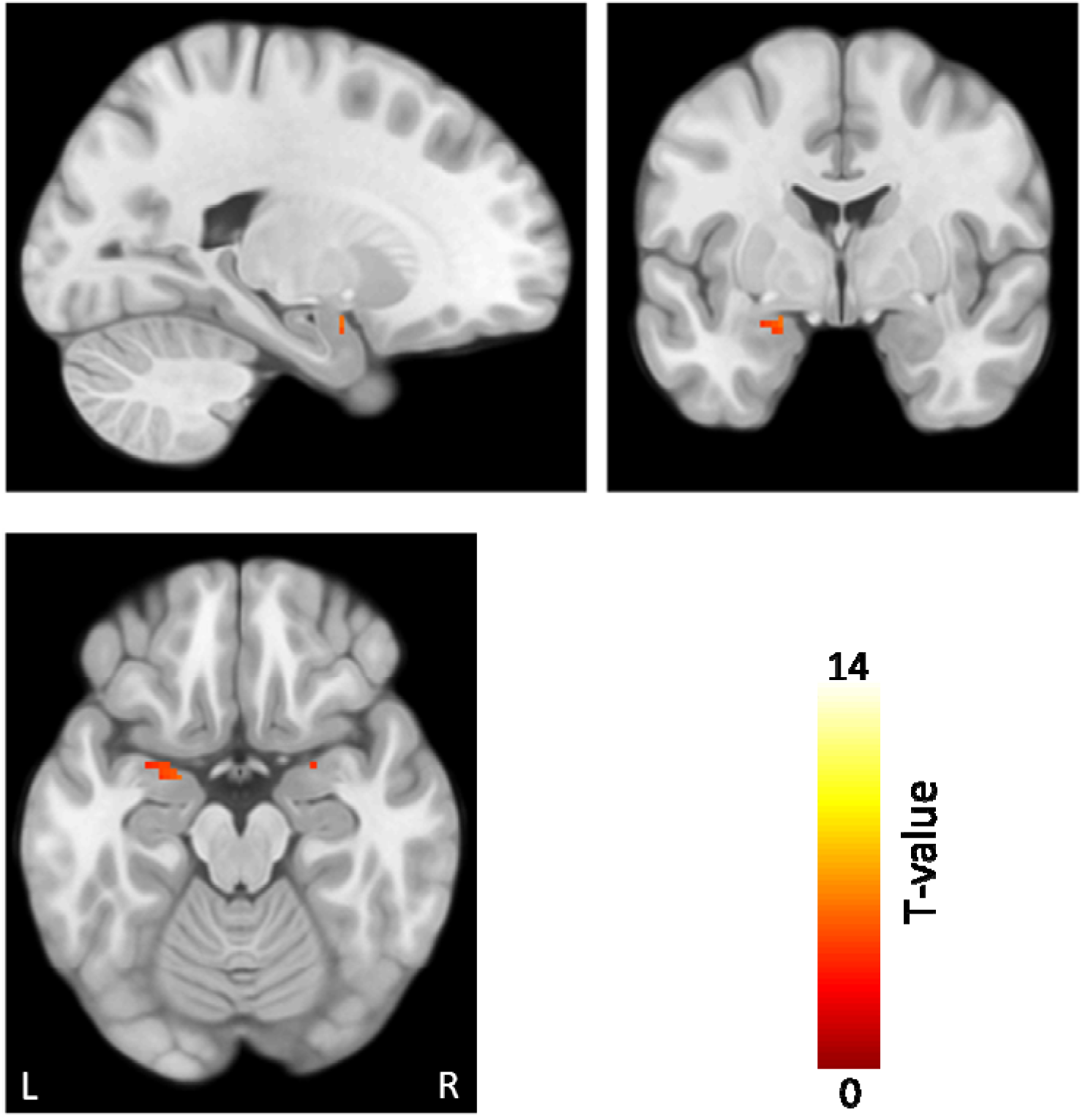
Medial temporal lobe responses to proportional increase in texture statistics in Experiment 1 (p< 0.05, FWE-corrected for medial temporal lobe volume). BOLD responses were measured while participants compared pairs of synthetic textures (a reference and a test) that varied in statistical level. Activity in the entorhinal cortex was selectively modulated by the level of texture statistics of the test sound.

### PPI Analysis

We performed PPI analysis to assess whether connectivity between auditory sub-regions and medial temporal lobe regions was modulated by texture naturalness in Experiment 1. The results revealed a significant negative modulatory effect of naturalness on the connectivity between the hippocampus (HC) and auditory sub-regions during the reference-sound presentation (Figure 5, Table 1) as well as within auditory sub-regions in the right hemisphere. Specifically, as the sound textures became more unnatural (i.e., lower statistical fidelity), functional connectivity increased between the HC and auditory areas (Te1.2 and Te2.1) in the left hemisphere, and between the HC and Te1.2, Te2.1, Te2.2, and Te3 in the right hemisphere (p < 0.05, FDR-corrected). This effect was evident in both seed configurations: significant connectivity was observed when the HC was used as the seed and when auditory subregions (Te1.2, Te2.1 and Te3) were used as seeds. This bidirectional pattern suggests that the functional connectivity between hippocampus and auditory regions is robust across different seed choices.

**Figure 5.**
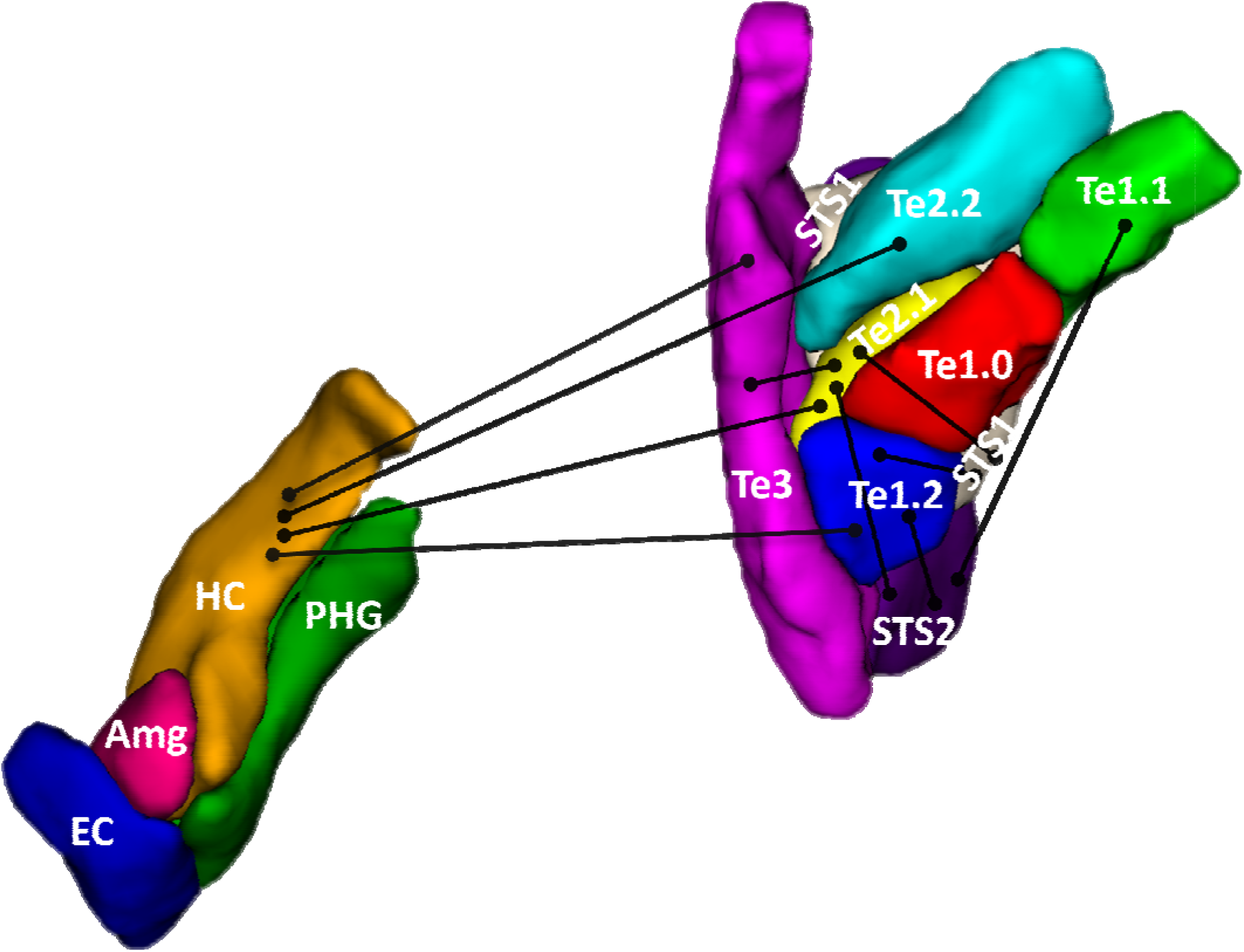
Group-level psychophysiological interaction (PPI) results showing connectivity between auditory regions and hippocampus modulated by sound texture statistics (p < 0.05, FDR-corrected) in Experiment 1. Results are displayed on a 3D reconstructed surface of the right hemisphere, highlighting anatomical regions of interest, visualised using ITK-SNAP (www.itksnap.org, Yushkevich et al., 2006). (HC = hippocampus, Amg = amygdala, EC = entorhinal cortex, PHG = parahippocampal gyrus, STS = superior temporal sulcus).

To determine whether sensory- HC connectivity was specific to the auditory cortex, a separate PPI analysis was conducted using visual regions (inferior, middle, and posterior occipital gyri) as seeds and MTL regions as targets, and vice versa. No significant modulatory effect of Δ on connectivity between visual and MTL areas was observed.

## Discussion

In this study, we investigated the neural representation of natural sound texture statistics. Across both experiments, auditory cortical responses increased with texture naturalness, indicating sensitivity to the statistical regularities that structure real-world sounds. Although primary and nonprimary auditory regions showed similar naturalness-dependent response patterns in both experiments, overall response magnitudes were lower in Experiment 2, where low-level acoustic features were held constant while naturalness was varied. This reduction suggests that low-level statistics contribute substantially to response strength, even though higher-order statistics alone remain sufficient to elicit graded responses. Together, these findings demonstrate that auditory cortex encodes sound texture statistics in a graded and distributed manner.

There is substantial evidence for a hierarchical organization of auditory processing for human speech and musical sounds. Nonprimary auditory regions tend to show category-selective responses, whereas primary regions predominantly encode low-level acoustic features such as spectral and temporal modulations (Leaver and Rauschecker, 2010; Okada et al., 2010; Moerel et al., 2012; Norman-Haignere et al., 2015; Norman-Haignere and McDermott, 2018; Landemard et al., 2021). In particular, sounds synthesized from spectrotemporal modulation-based models often evoke strong responses in primary auditory cortex but weaker responses in nonprimary regions, implying that nonprimary areas are sensitive to higher-order features present in natural speech and music that such models fail to capture (Norman-Haignere and McDermott, 2018). Indeed, McDermott and Simoncelli (2011) showed that participants rated synthesized speech and music as highly unrealistic, even when the synthetic sounds were statistically matched to the original recordings. This indicates that the perception of speech- and music-specific attributes relies on features or computations not represented in the auditory model.

In contrast, synthesized sound textures generated from auditory models are perceptually similar to their real-world counterparts highlighting that theses texture statistics might be sufficient for recognition (McDermott and Simoncelli, 2011). In our study, we used synthetic stimuli in which statistics were parametrically varied, producing sounds that ranged from least natural to natural. Across both experiments, we observed a similar pattern of graded responses distributed across primary and nonprimary auditory regions, but weaker responses were observed in the second experiment. Based on prior work, one might expect that controlling low-level acoustic cues while varying naturalness would primarily reduce responses in primary auditory cortex, with higher-order regions remaining sensitive to higher-level texture statistics. However, the observed reduction in nonprimary regions (e.g., Te3 and neighbouring areas) suggests that the hierarchical organization commonly reported for speech and music, where these regions show sensitivity only for high-level features, may not generalize as clearly to natural sound textures. Instead, our findings indicate a more distributed representation of texture naturalness across the auditory cortex.

When the sounds were matched for modulation power, we observed reduced discriminability between the stimuli with the largest naturalness difference, as well as weaker responses across most auditory regions (with the exception of Te1.2 and Te2.2) in Experiment 2. This pattern suggests that holding modulation power constant reduced the effective range of naturalness variation, making even the largest naturalness contrasts less perceptually distinguishable than in Experiment 1. The corresponding reduction in neural responses further supports the auditory cortex’s sensitivity to texture naturalness and aligns with previous findings showing that modulation power is a driver of cortical responses to complex natural sounds (Theunissen et al., 2000; Overath et al., 2015). Crucially, the presence of significant neural responses in Experiment 2, even with fixed modulation power, indicates that the effects observed in Experiment 1 cannot be attributed solely to differences in this statistic. Instead, the results suggest that multiple statistical components contribute differentially to overall response magnitude, while the underlying representational architecture remains distributed across auditory cortex.

Animal studies using natural textures, however, report stronger sensitivity to high-order statistics in the inferior colliculus (IC) than in auditory cortex, and show that texture structure is reflected in neural spectrum and neural-correlation statistics of IC ensembles, which subserve recognition and discrimination roles (Peng et al., 2024). We did not observed responses in IC to change in texture naturalness. Nonetheless, our findings reveal graded responses in nonprimary auditory areas such as Te3, an area repeatedly shown to be selective for human speech and music (Leaver and Rauschecker, 2010; Okada et al., 2010; Moerel et al., 2012; Norman-Haignere et al., 2015; Norman-Haignere and McDermott, 2018; Landemard et al., 2021) as well as in primary regions.

In vision, studies by Freeman et al. (2013) and Ziemba et al. (2016, 2024) demonstrated differential neuronal responses in areas V1 and V2 of anesthetized macaques and in humans when viewing naturalistic texture images that preserve higher-order statistical dependencies, compared to spectrally matched noise. While neurons in V1 responded similarly to both stimulus types, V2 neurons exhibited significantly stronger responses to the naturalistic textures. These findings suggest that cortical sensitivity to texture statistics in vision is organized hierarchically. In contrast, as noted above, our results indicate that a similarly pronounced hierarchical pattern may not be as evident in the auditory system for natural sound textures. After controlling for low-level acoustic features, we observed decreased responses in both primary and nonprimary auditory regions as sound textures became less natural. This pattern suggests that sensitivity to higher-order statistical structure in sound textures is more broadly distributed across auditory cortex, rather than reflecting a clear hierarchical transition comparable to that observed in vision.

In Experiment 1, in addition to the auditory cortical effects, we found that activity in the entorhinal cortex (EC) increased in proportion to the texture statistics present in the test sound. Our task required participants to encode the reference (the first sound) texture and then compare it to the test sound (the second sound), which likely engaged networks beyond those involved in perception alone. The fact that EC activity was modulated by texture statistics during the comparison phase (the test sound) suggests that the EC may contribute to perceptual comparison processes rather than only memory retrieval. We did not observe a similar effect in Experiment 2, where modulation power was controlled; this absence may reflect reduced statistical power due to the smaller sample size. Overall, these results point to an effect driven by task demands rather than a central role of medial temporal lobe regions in representing texture statistics.

Connectivity analyses further revealed stronger interactions between auditory cortex and the hippocampus, as well as between auditory subregions and the superior temporal sulcus (STS), when the reference textures were unnatural (i.e., lower Δ). This effect could reflect top-down perceptual inference under uncertainty (Asilador and Llano, 2021) or enhanced memory encoding encoding (Kumar et al., 2016; Dimakopoulos et al., 2022), given that participants were required to maintain the reference sound in memory during the delay period. The increased connectivity between auditory cortex and STS was unexpected and may indicate elevated processing demands when textures were less natural. Overall, these findings contribute to growing evidence for functional interactions between the hippocampus and the auditory system (Billig et al., 2022; Zhang et al., 2022; Leong et al., 2023) and provide new insight into auditory–STS connectivity.

## Conclusion

These findings demonstrate that the auditory cortex shows a graded and distributed sensitivity to natural sound texture statistics. BOLD responses in both primary and nonprimary auditory regions increased with texture naturalness; however, when modulation power was controlled, overall response magnitudes were reduced. This reduction suggests that low-level statistics contribute substantially to neural response strength, even though higher-order statistics alone remain sufficient to elicit a graded response. Medial temporal lobe regions were engaged primarily under conditions of greater perceptual uncertainty or increased task demands, consistent with a modulatory rather than representational role in texture processing.

## Supporting information

Figure S1. Texture Cochleagram and summary statistics. All statistics (cochlear channel variance, kurtosis and skewness, cochlear correlation, modulat

## Conflict of interest statement

The authors declare no competing financial interests.

## Acknowledgments

This work was supported by the Medical Research Council, United Kingdom (MR/T032553/1). We thank Martina Callaghan, Fred Dick and Oliver Josephs for designing the sound delivery system and MRI sequences; Karl Friston and Peter Zeidman for their advice on experimental design; Richard McWalter for providing code and commenting on the manuscript; and the Imaging Support team at the UCL Department of Imaging Neuroscience.

## Notes

https://osf.io/rvena/?view_only=2d7ee5ca4f0c4b32a1239ca56ba3a857

### Competing Interest Statement

The authors have declared no competing interest.

### Summary of Updates

A new experiment has been done; Figure 1 added; Supplemental files updated.

