## Supplementary figures and images for "Brain Representations of Natural Sound Statistics"

### Figure S1. Texture Cochleagram and summary statistics. All statistics (cochlear channel variance, kurtosis and skewness, cochlear correlation, modulat

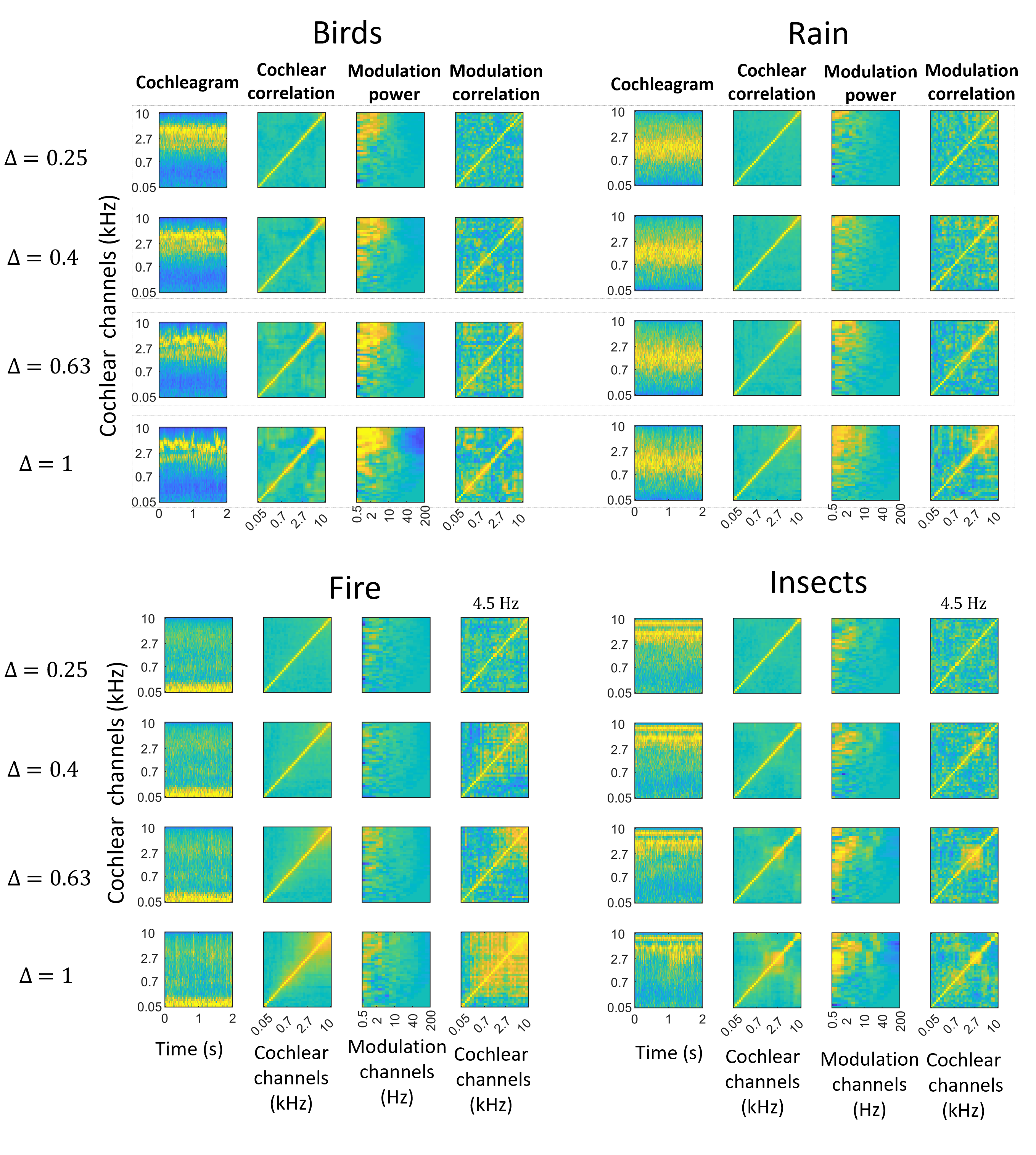

### Supplemental Data 1

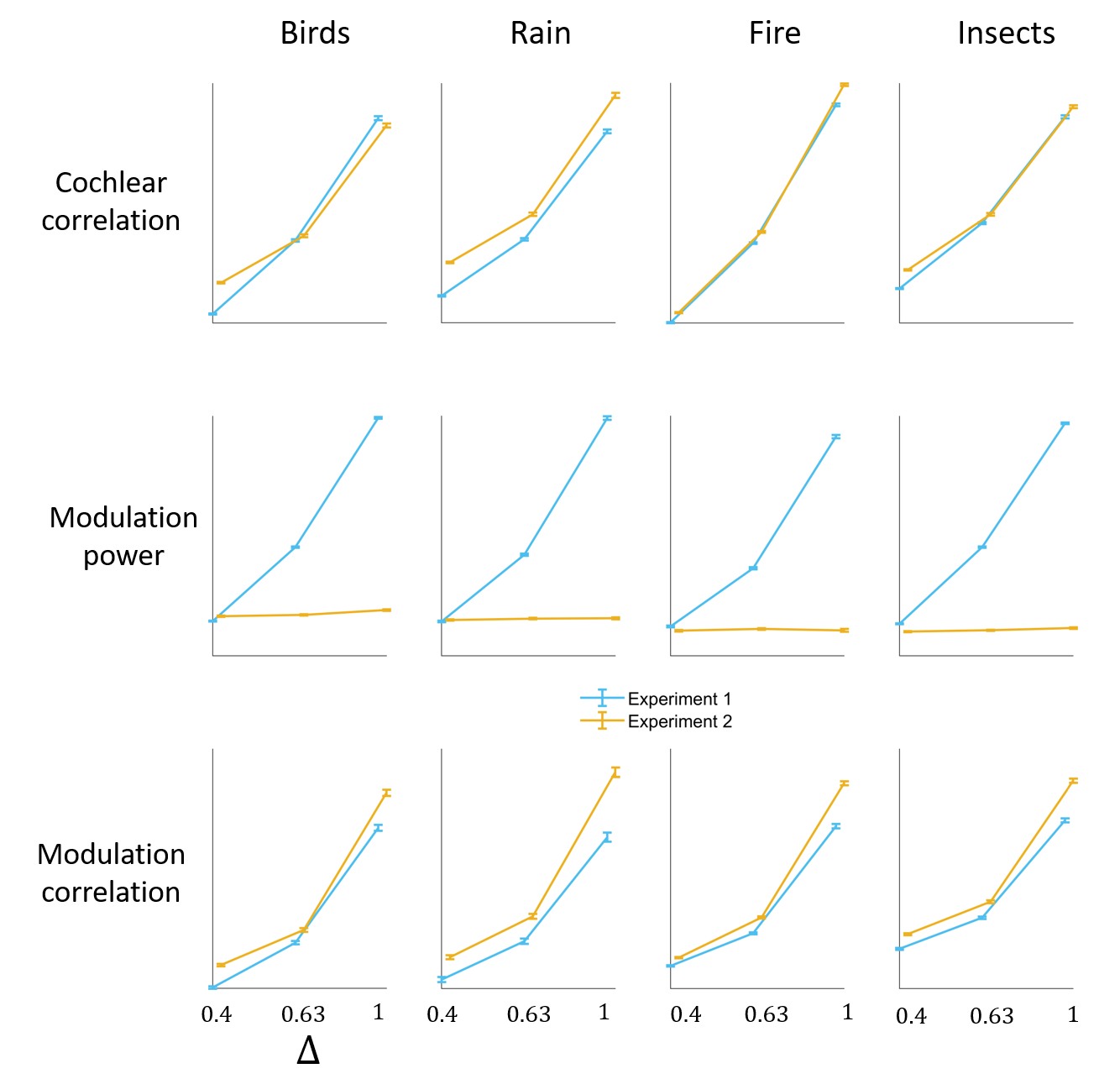
